# Detecting haplotype-specific transcript variation in long reads with FLAIR2

**DOI:** 10.1101/2023.06.09.544396

**Authors:** Alison D. Tang, Eva Hrabeta-Robinson, Roger Volden, Christopher Vollmers, Angela N. Brooks

## Abstract

**Background:** RNA-Seq has brought forth significant discoveries regarding aberrations in RNA processing, implicating these RNA variants in a variety of diseases. Aberrant splicing and single nucleotide variants in RNA have been demonstrated to alter transcript stability, localization, and function. In particular, the upregulation of ADAR, an enzyme which mediates adenosine-to-inosine editing, has been previously linked to an increase in the invasiveness of lung ADC cells and associated with splicing regulation. Despite the functional importance of studying splicing and SNVs, short read RNA-Seq has limited the community’s ability to interrogate both forms of RNA variation simultaneously.

**Results:** We employed long-read technology to obtain full-length transcript sequences, elucidating cis-effects of variants on splicing changes at a single molecule level. We have developed a computational workflow that augments FLAIR, a tool that calls isoform models expressed in long-read data, to integrate RNA variant calls with the associated isoforms that bear them. We generated nanopore data with high sequence accuracy of H1975 lung adenocarcinoma cells with and without knockdown of *ADAR*. We applied our workflow to identify key inosine-isoform associations to help clarify the prominence of ADAR in tumorigenesis.

**Conclusions:** Ultimately, we find that a long-read approach provides valuable insight toward characterizing the relationship between RNA variants and splicing patterns.

**Highlights:**

- FLAIR2 has improved transcript isoform detection and incorporates sequence variants for haplotype-specific transcript detection.
- In addition to haplotype-specific variant detection, it identifies transcript-specific RNA editing
- Able to identify haplotype-specific transcript isoform bias in expression
- Long-read sequencing identifies hyperedited transcripts that are missed from short-read sequencing methods for a more comprehensive identification of ADAR targets

## Introduction

Adenosine-to-inosine (A-to-I) editing is one of the most common forms of RNA editing in organisms with a developed central nervous system [1–5]. As inosines are recognized by cellular machinery as a guanosine, one potential downstream effect of A-to-I editing is the alteration of coding sequence. There are numerous cases of A-to-I recoding identified as essential for normal brain function [6–8] and yet other cases where recoding worsens disease prognosis [9–11]. In addition to recoding potential, inosines can affect RNA splicing in a *cis-*regulatory manner through the disruption of splice sites or splicing regulatory elements, leading to the creation of alternatively spliced mRNAs [12–14]. Considering that 95-100% of multi-exon genes are alternatively spliced [15], the effects of A-to-I editing on coding changes, regulatory elements, and alternative splicing require further study to elucidate.

The expression of ADAR1 is ubiquitous and the A-to-I editing that ADAR1 mediates on dsRNAs is widespread [16]. Previous literature has described the role of ADAR1 in autoimmune diseases [17–19], such as in the case of decreases in editing levels in particular dsRNAs resulting in MDA5-dependent interferon response and inflammation [20]. Additionally, aberrant ADAR activity has been linked to many other diseases [6,7,21–25], including diseases of the lung and blood, in which ADAR overexpression is associated with increased malignancy [10,11]. In H1975 lung adenocarcinoma (ADC) cell lines, ADAR is not only upregulated but also has been shown to bind to and edit focal adhesion kinase (FAK), increasing both FAK expression and mesenchymal properties of the cells [10]. The connection of ADARs with diseases, in particular lung adenocarcinoma, in addition to the influence that ADARs have on the transcriptome underscores the importance of characterizing the complete RNA sequences that bear inosine edits.

Despite appreciable efforts to map A-to-I editing sites [4,26], there is an absence of studies examining the full transcriptional context of inosines. Previous efforts to document A-to-I editing using short-read sequencing report the genomic position of edited sites [4,26,27] and our goal is to analyze the transcripts where edits reside. To investigate the transcriptome-wide impact of ADAR in lung ADC, we performed nanopore long-read cDNA sequencing of H1975 lung ADC cells with ADAR knockdown. We overcame the relatively high error rate of nanopore sequencing by using the Rolling Circle Amplification to Concatemeric Consensus (R2C2) nanopore cDNA sequencing method [28]. R2C2 greatly lowers the error rate of nanopore cDNA sequencing through the increase of single molecule coverage, yielding a median 98.7% base accuracy [29]. Accurate, long reads allow us to resolve full-length transcripts and RNA editing, equipping us to better understand the role of ADAR editing in the cancer transcriptome.

In RNA-seq data, there is ambiguity as to whether mismatches to the reference genome correspond to 1) somatic or germline variants, 2) RNA edits in which one nucleotide is edited to read as another, or, in the case of nanopore direct RNA sequencing, 3) modified RNA nucleotides. Although R2C2 is unable to preserve RNA modifications, we have devised a tool to phase and associate consistent mismatches to isoform models given long reads, agnostic to the kind of alteration that accounts for the mismatch. We refer to these mismatch-aware isoforms generally as haplotype-specific transcripts (HSTs), with a set of variants occurring on the same transcripts designated a “haplotype”. In efforts to jointly identify isoform structure and the potentially stochastic nature of inosine positions in nanopore data, we introduce a computational software for identifying HSTs. We built upon the isoform detection tool FLAIR [30], which is one amongst many tools (Stringtie2 [31], FLAMES [32], TALON [33], MandalorION [34]) developed for this purpose. FLAIR was initially developed to identify transcript models in long reads to hone in on subtle splice site changes; the original FLAIR method was primarily concerned with error-prone and truncated reads, with minimal consideration for sequence variation. This variant-aware FLAIR, called FLAIR2, incorporates mismatches from a variant caller into transcript models for an arbitrary number of haplotypes as would be useful for grouping editing events, distinguishing itself from other allele-specific expression tools for long reads such as LORALS [35], IDP-ASE [36], and FLAMES.

Here, we use FLAIR2 to detect haplotype-specific transcripts in a diploid mouse hybrid long and short read dataset and compare changes in inosine editing in the context of lung cancer. We sequenced lung ADC replicates with *ADAR1* knocked down using Illumina RNA-Seq as well as R2C2 nanopore sequencing. Paired with the development of the necessary computational framework for full-length isoform and RNA editing analyses, we reveal new insights into long-range A-to-I edits and demonstrate the power of nanopore sequencing as a tool for the transcriptome-wide identification of inosines.

## Results

### FLAIR2 is a variant-aware isoform detection pipeline

In an effort to build user-friendly computational workflows for nanopore data, we previously developed a tool called Full-Length Alternative Isoform analysis of RNA (FLAIR). FLAIR calls isoform structures and performs various isoform-level analyses of nanopore cDNA and direct RNA sequencing data [37]. We designed the FLAIR workflow to account for the increased error rate of long reads, in particular for nanopore data. Previous work with FLAIR emphasized the discovery of isoform models and their comparison between sample conditions. We have adjusted FLAIR to incorporate phased variant calls to investigate haplotype-specific transcript expression in nanopore data. We also sought to improve FLAIR’s performance on isoform structure (transcript start and ends and exon-exon connectivity) by increasing sensitivity to annotated transcript isoforms. Here, we evaluated FLAIR2 performance on isoform structure using SIRVs [38]; a more comprehensive evaluation of FLAIR2 is being performed through the Long-read RNA-seq Genome Annotation Assessment Project (LRGASP) Consortium [39][40].

The modified FLAIR workflow (FLAIR2) now begins with an alignment of all reads to the annotated transcriptome. The addition of this ungapped alignment step was designed to improve small or microexon detection for error-containing, spliced reads which are difficult to align to the genome [41]. Reads are assigned to an annotated transcript if they have high sequence identity with the transcript, with an emphasis of accuracy proximal to splice sites (see “FLAIR splice site fidelity checking” in Methods). The annotated transcripts that have sufficient long read support are included as part of the set of FLAIR isoforms. The remaining reads that are not able to be assigned to an annotated transcript are then used to detect novel transcript models (see Methods). The final, sample-specific isoform assembly includes the supported, annotated isoform models combined with the novel models. FLAIR is also capable of downstream analyses such as isoform quantification and differential expression tests of nanopore data, as described previously [30].

We compared the performance of FLAIR2’s updated isoform detection method with Stringtie2 [31] and FLAMES [32] on SIRVs sequenced with nanopore R2C2 sequencing (see Methods). Transcript detection with FLAIR2 has a higher precision of 95.5 (Table 1), indicating fewer false positive transcripts detected compared to other tools, with comparable sensitivity with the other tools. We also investigated the transcript-level precision and sensitivity using nanopore 1D cDNA SIRV sequences, where FLAIR2 again performed best comparatively in precision and performed similarly to other tools in terms of sensitivity (Table 2). On these SIRVs, FLAIR2 demonstrated marked improvement (37 point increase in transcript-level precision) over the previously published FLAIR, which focused more on nucleotide-level sensitivity and precision.

**Table 1.** Performance of transcript detection tools on R2C2 nanopore SIRVs.

|  | Transcript-level sensitivity | Transcript-level precision |
| --- | --- | --- |
| FLAIR2 | 77.8 | **95.5** |
| FLAMES | **81.5** | 94.3 |
| Stringtie2 | 72.8 | 90.8 |

**Table 2.** Performance of transcript detection tools on 1D nanopore SIRVs.

|  | Transcript-level sensitivity | Transcript-level precision |
| --- | --- | --- |
| FLAIR2 | 63.8 | **89.8** |
| Stringtie2 | **76.8** | 81.5 |
| FLAMES | 66.7 | 78.0 |
| FLAIR | 65.1 | 51.9 |

### Assessing FLAIR2 for haplotype-specific transcript detection

To be able to integrate sequence variants into transcripts detected with FLAIR, we applied both longshot [42] and PEPPER-Margin-DeepVariant [43] to call variants in long-read data. Both variant callers were developed for diploid variant calling and phasing in long reads. Following isoform identification, FLAIR has two modalities for phasing variants to discover variant-aware transcript models. The first uses phasing information from longshot, which is comprised of a phase set determined for each read and a set of variants corresponding to each phase set. For the second modality, as we anticipated working with RNA edits and potential cancer-related aneuploidies that may result in more than two consistent haplotypes, FLAIR can approach phasing variants that is agnostic to ploidy: 1) given variant calls, FLAIR tabulates the most frequent combinations of variants present in each isoform from the supporting read sequences; 2) from the isoform-defining collapse step, FLAIR generates a set of reads assigned to each isoform; so 3) isoforms that have sufficient read support for a collection of mismatches are determined (Fig 1a). This latter method of phasing focuses solely on frequency of groups of mismatches that co-occur within reads and does not use ploidy information to refine haplotypes, allowing for the generation of multiple haplotypes within a gene and transcript model.

**Figure 1.**
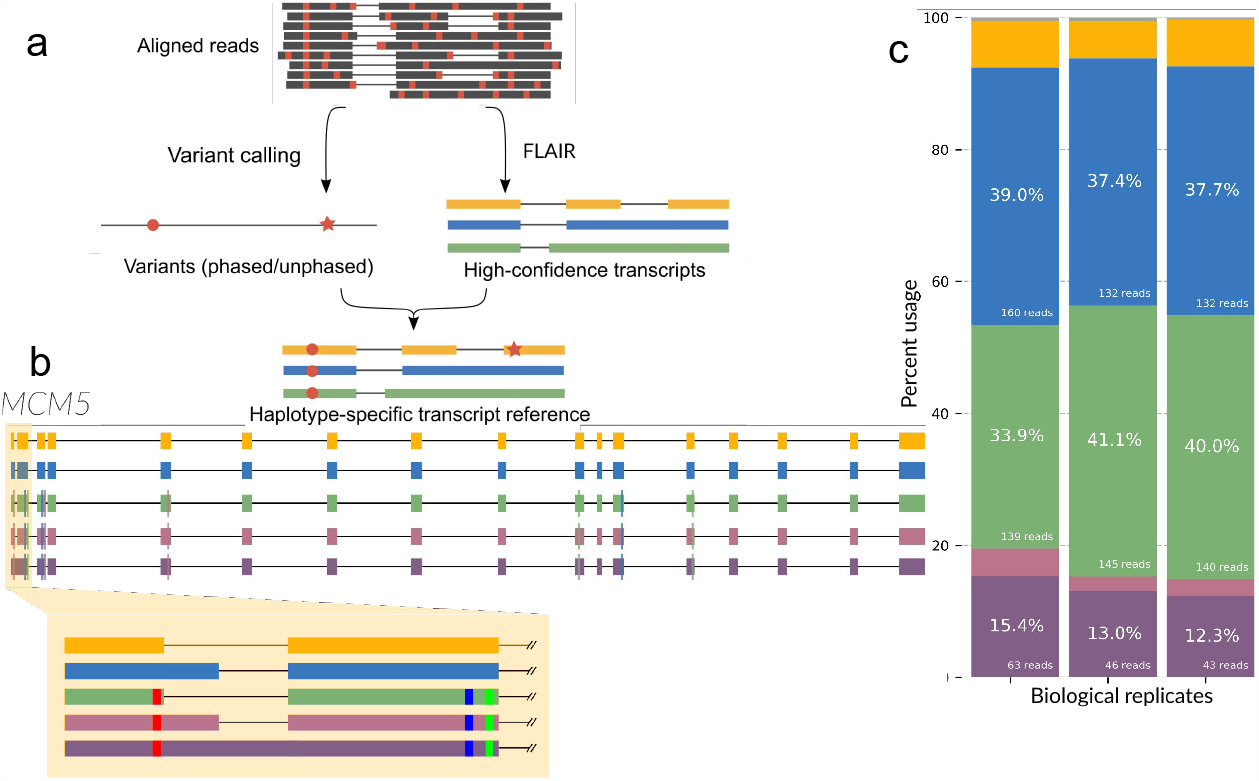
Variant-aware transcript detection by FLAIR2 identifies haplotype-specific transcript isoform bias. a, FLAIR2 computational workflow for identifying haplotype-specific transcripts in long reads. **b**, FLAIR transcript models for *MCM5* with the highest expression are plotted using different colors for each transcript’s exons. The highlighted portion shows alternative splicing and the smaller blocks within exons indicate variants. **c**, Stacked bar chart showing the proportion of transcript expression of transcripts from **b** as matched by color for each of the replicates sequenced.

We tested the FLAIR2 isoform discovery pipeline on R2C2 data generated from Castaneus x Mouse 129 hybrid mouse embryonic stem cells[39] where we expect evidence of HSTs partitioned by known parental haplotypes. Integrating longshot’s phased diploid variant calls, we identified transcripts that were significantly associated with a haplotype compared to other transcripts in the gene (Fisher’s exact adjusted p-value < 0.05) and then analyzed the set of 152 genes that contained HST bias. Nine of these genes were known imprinting genes [44]. GO analysis of HST-containing genes reveals an enrichment in DNA repair and damage terms (Supplementary Table 1,2), an attribute of embryonic stem cells for maintaining genomic integrity [45,46]. One example from these gene sets is *MCM5*, a component involved in the DNA helicase complex [47]. The non-reference haplotype that longshot reports corresponds to the *castaneus* parent haplotype [48] and exhibits HST bias (Fig 1b,c). The haplotype, which contains a variant close to the 5’ splice site of the first exon, is coupled with either splicing with the proximal 5’ splice site or a retained intron. With these results in mice, we are able to demonstrate FLAIR2’s capacity for incorporating variants in diploid transcriptomes and detecting HSTs in long reads.

### Global downregulation of A-to-I editing following ADAR1 knockdown in short and long reads

We applied FLAIR2 to study isoform alterations in relation to inosine editing to build on our understanding of A-to-I editing in the cancer transcriptome. Previous work [10] discovered a connection between A-to-I editing, FAK (PTK2) transcript stability, and increased malignancy using short-read sequencing. We followed their approach of knocking down ADAR and investigating alterations in editing, however we leveraged the combination of long- and short-read cDNA sequencing (Fig 2a) to resolve the full-length transcripts edited in lung ADC. First, ADAR1 knockdown was performed in H1975 cells using three different ADAR1 siRNAs (see Methods) to achieve 70-80% knockdown of ADAR1 protein levels compared to control replicates (Fig 2b). Next, we sequenced the cDNA from three ADAR knockdown and three control knockdown samples. We observed a 55.1, 73.7, and 78.8% decrease in ADAR expression from our normalized Illumina RNA-Seq replicates, with ADAR being the most significantly downregulated gene (Supplementary Table 3, Fig 2c). We prepared barcoded R2C2 cDNA libraries for each of our samples and sequenced them in matched ADAR and control knockdown pairs using three MinIONs. We obtained an average of 11.7 gigabases with median read length 9,599 bp from each MinION (Table 3). We report a median accuracy of 99.3% and median read length of 1,287 bp from our consensus-called reads. As the number of consensus called and demultiplexed reads provided less power in separate replicates, we decided to pool all of the samples in each condition together for further analyses.

**Figure 2.**
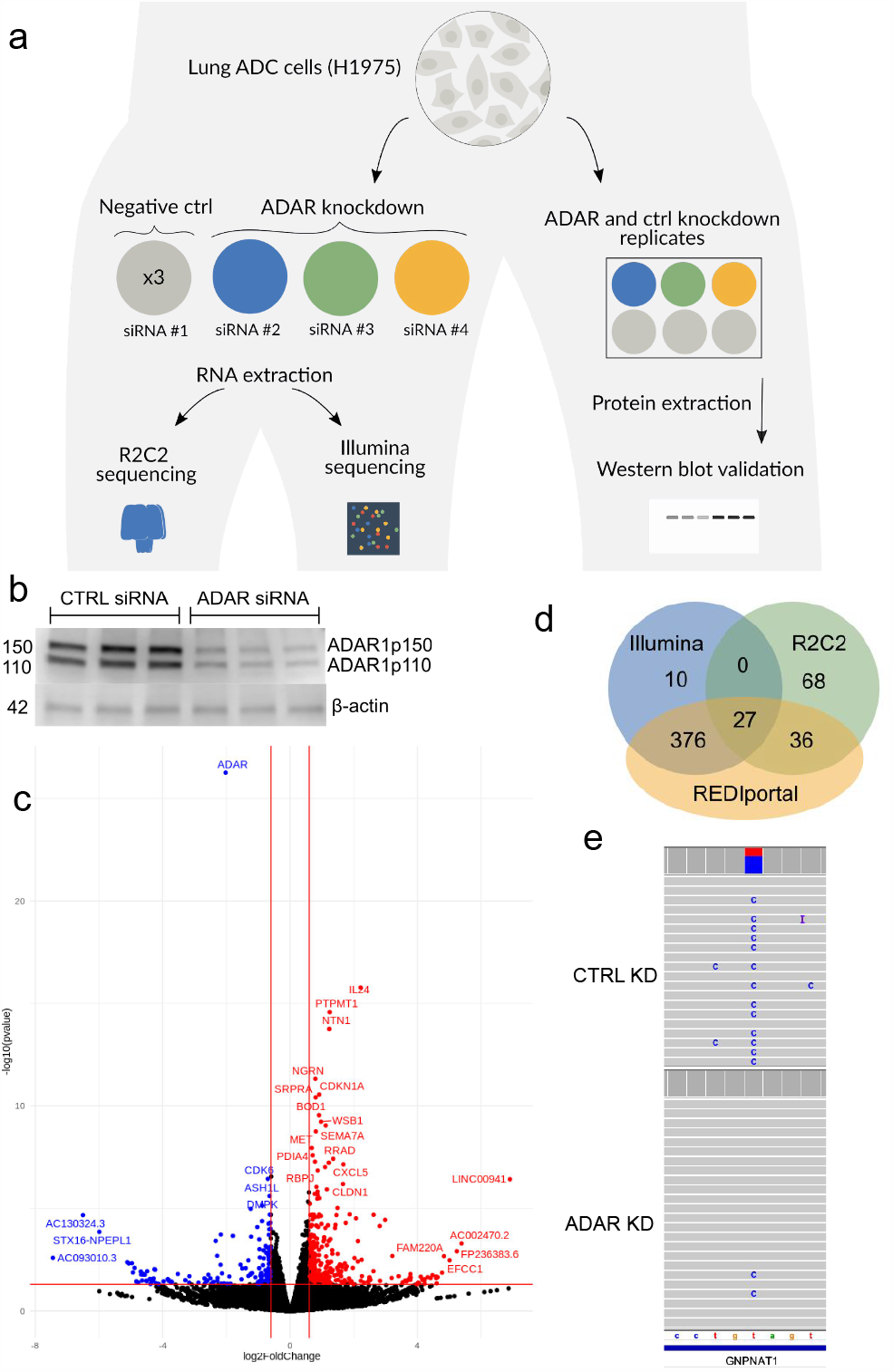
Identification of downregulated inosines with short- and long-read RNA-Seq. **a**, Experimental workflow of ADAR knockdown in H1975 cells **b**, Western blot validation of ADAR knockdown. **c**, Volcano plot of differentially expressed genes identified from Illumina sequencing. Red: genes with increased expressed after ADAR knockdown; blue: genes whose expression went down; black: no change in expression. **d**, Venn diagram comparison of the significantly downregulated inosines identified with Illumina, R2C2 nanopore, or present in the REDIportal database (hg38 liftover). **e**, IGV browser view of a downregulated inosine at chr14:52775760 in *GNPNAT1* in the R2C2 data.

**Table 3.**
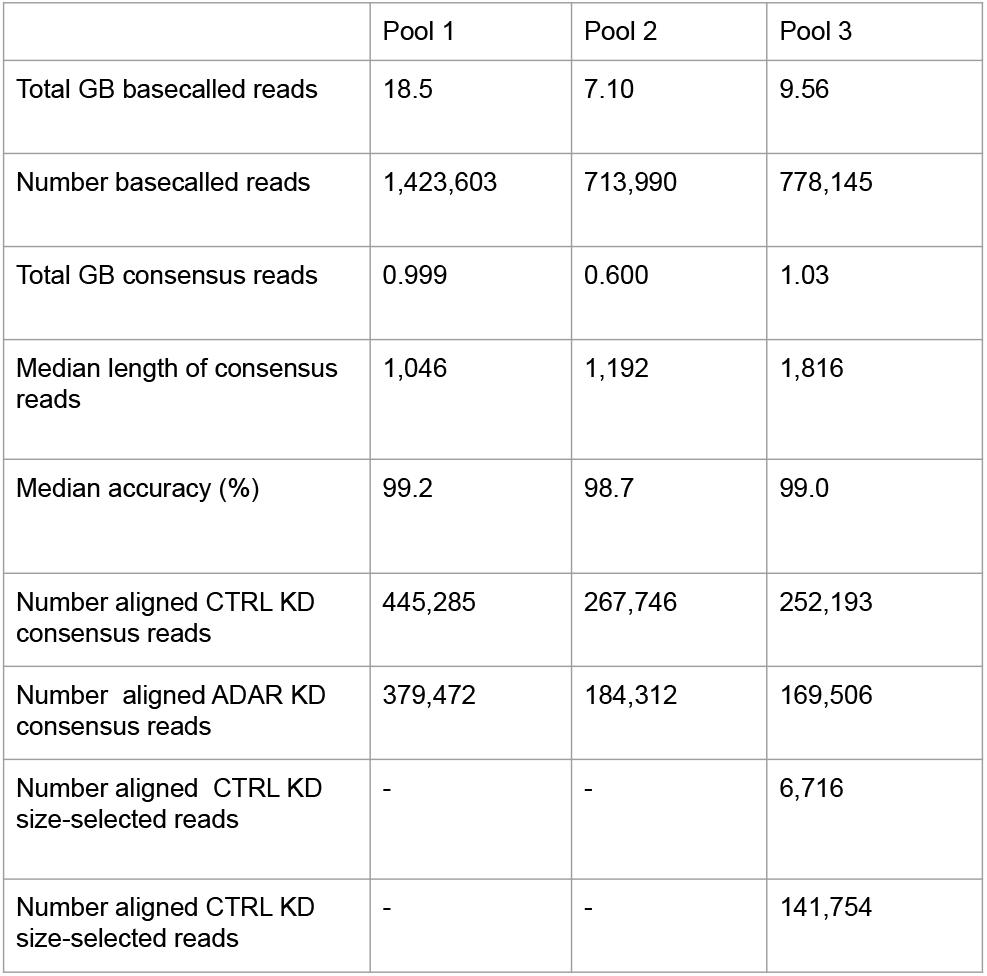
R2C2 nanopore sequencing numbers. For each ADAR KD and control KD sample pool that was sequenced on a MinION, we report the total number of reads obtained from sequencing after basecalling, consensus calling, and minimap2 alignment to the hg38 genome. We also show the number of gigabases of reads after basecalling and consensus calling, as well as the median length of the consensus reads. We calculated an accuracy for each read and report the median for each sequencing run.

### Inosine detection in short and long reads

We used reditools [49] to catalog nucleotides at every position in the Illumina data and filtered for the positions that conformed to A-to-I expectations (i.e. positions with an A or T in the reference and read support for G or C). We identified 334 A→G mismatches in the Illumina data that were significantly changed upon ADAR knockdown (Methods), with the majority (324) of these positions present in the REDIportal database [49]. Of these 334 A-to-I events, 312 were downregulated in the knockdown conditions and 12 were upregulated.

We considered longshot and PEPPER-Margin-DeepVariant variant calls to identify an initial set of A-to-G mismatches that we would then reclassify as A-to-I edits with REDIportal and downregulation analyses. Both variant callers identified variants that could be categorized as inosine changes that the other caller missed; as such, we combined all the variant calls from both tools for increased sensitivity. Starting with the combined variant calls, we identified 63 significantly changed A-to-I events that were also present in REDIportal (Fisher’s exact p<0.05) with a greater than 10% difference in proportion of edited reads (Fig 2d, Supplementary Data File 1). As expected, most (62/63) were downregulated in the ADAR knockdown samples. Of the 131 significant nanopore-identified inosines, 27 were also identified as significantly downregulated in the Illumina data (Fig 2d,e). We identified individual bases with a high proportion of editing, defined as type I hyperediting following nomenclature from a previous study which considered bases with >40% of adenosine residues being edited as type I hyperediting [50]. We found that approximately half (79/131) of the significantly downregulated inosines were considered type I hyperedited in the control knockdown data. Some inosines identified as significantly differentially edited with nanopore but not in Illumina were events that received insufficient aligned coverage or did not have enough edited reads to pass a significance threshold (Fig 3, Supplementary Fig 1). In conclusion, while the coverage of short read data will typically surpass that of long reads lending to an increase in the number of inosines detected, long reads are advantageous for detecting certain novel A-to-I events.

**Figure 3.**
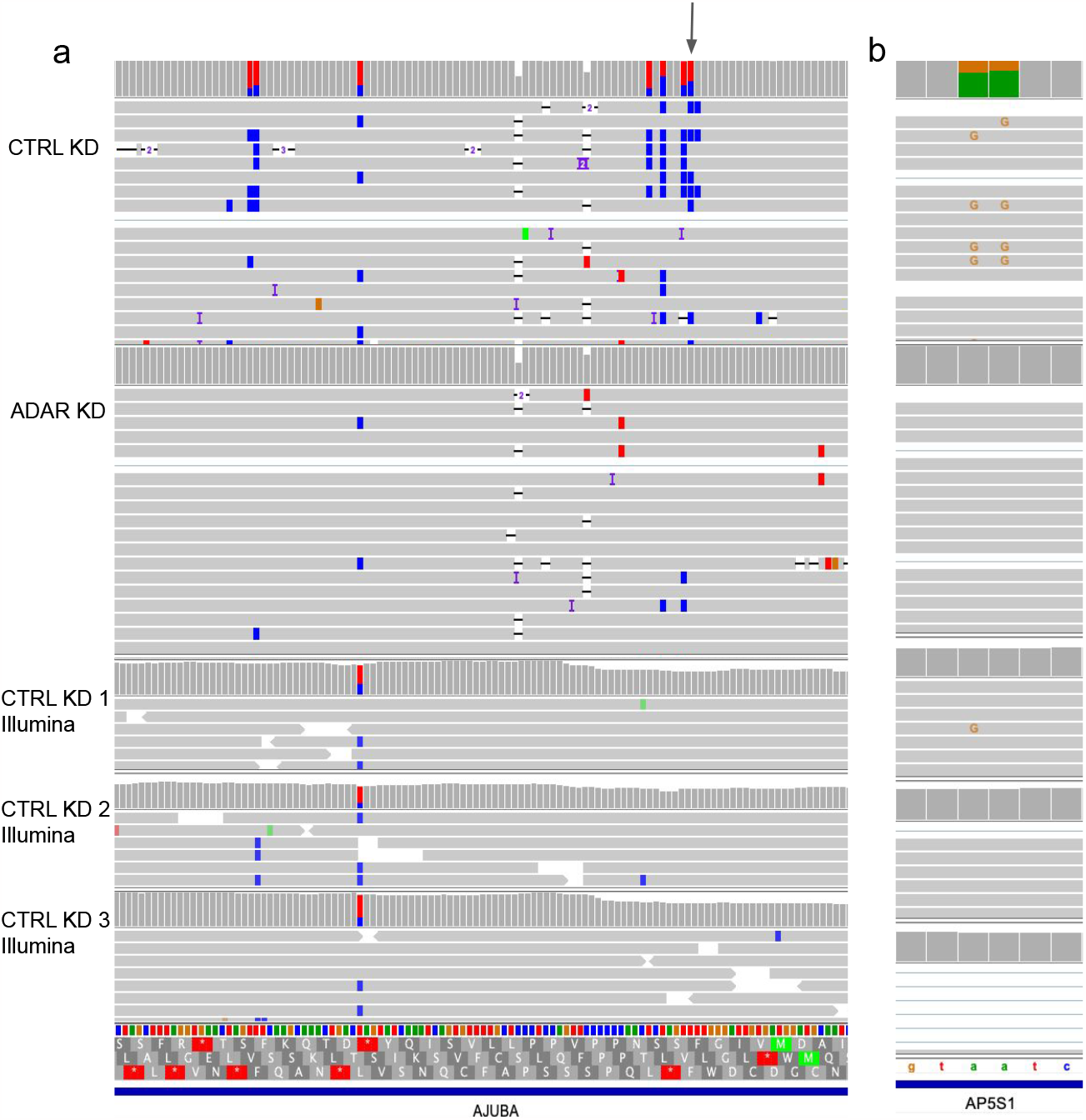
Significantly downregulated A-to-I detected with nanopore and not in the Illumina data. IGV shots of nanopore and Illumina data aligned to hg38 **a**, Gray arrow indicates the differentially edited position found in nanopore but not Illumina and is a known editing position in REDIportal. **b**, Known A-to-I editing detected by nanopore reads in *AP5S1* but missed in Illumina.

### Long reads can identify type II hyperediting

ADAR tends to produce clusters of inosines on a transcript, which we define as type II hyperediting [50]. Hyperedited regions were identified as any window that contained at least three A-to-I edits distributed within every 150 bp—a more relaxed definition derived from Porath et al. [56] which requires >5% of a read’s length to contain A-to-G mismatches. Type II hyperedited transcripts have been associated with nuclear retention or degradation [51–55]. First, we note a pattern of ADAR editing in which transcripts that are edited tend to have multiple edits. The control knockdown data in aggregate show that 38.7% of reads contain at least one edit, and of the reads that are edited, 77.9% contain more than one edit. On detecting multiple edits in short-read RNA-Seq, if the edits are too distant, or if a read contains many mismatches on account of A-to-I hyperediting (type II), reads with multiple edits may not align to the genome and evade detection [56] (Fig 3, Supplementary Fig 1). To expand our search space, we used the larger set of all inosines found in our nanopore data and REDIportal that were not necessarily significantly downregulated after knockdown as well as the novel significantly downregulated inosines discovered with nanopore only. With this approach, we identified 99 regions that overlapped with known type II hyperediting [56] as well as 17 novel hyperedited regions (Fig 4a).

**Figure 4.**
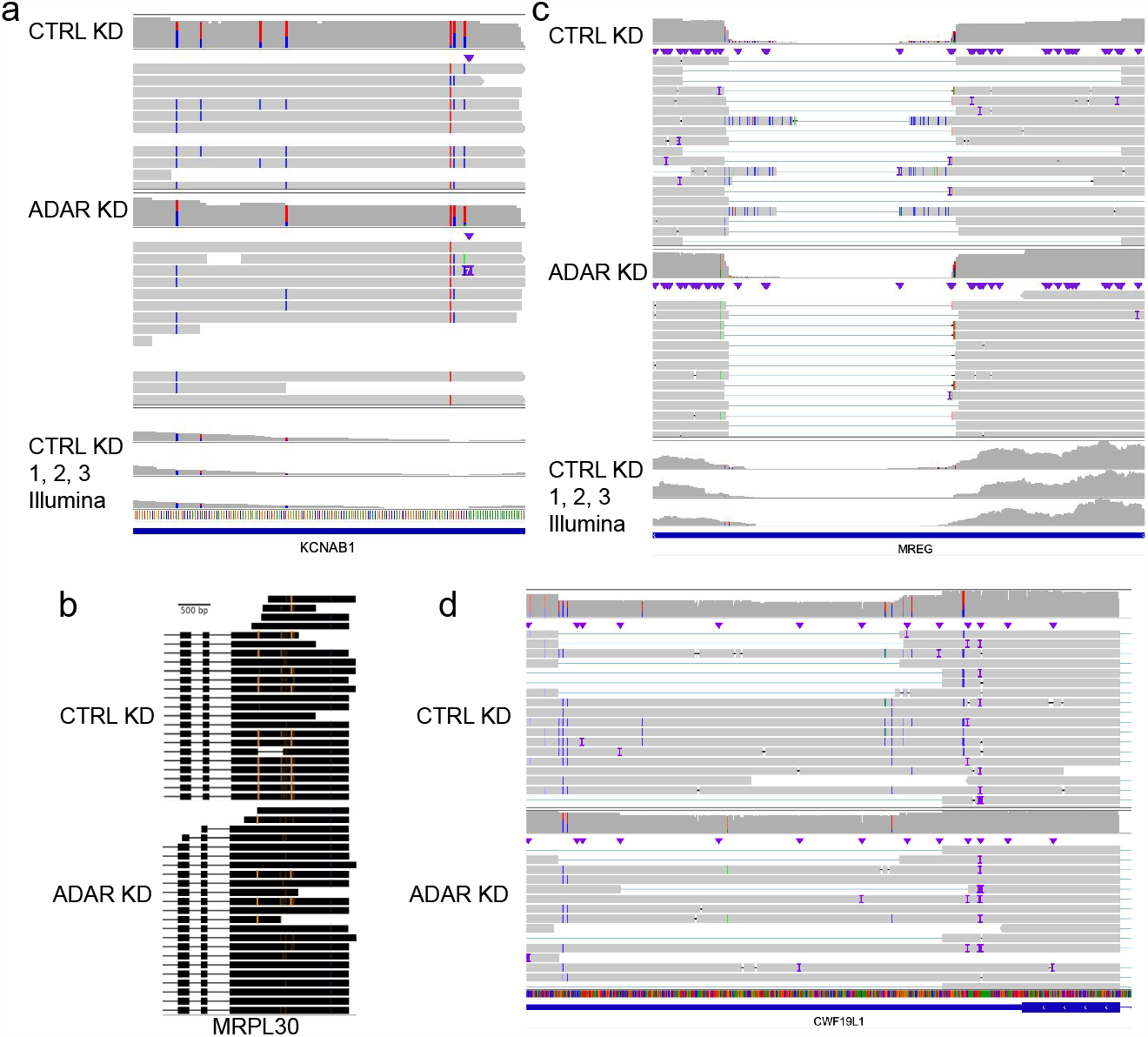
Long-range features of inosines observed with nanopore sequencing. Aligned reads displaying **a** type II hyperediting, **b** coordinated editing, and **c** and **d** disruption of splicing in the presence of editing.In **a** and **c**, the top three coverage tracks are Illumina CTRL KD samples and the bottom coverage track and reads are displaying the nanopore CTRL KD reads. In **b** and **d**, the dataset on top displays the control nanopore reads and the bottom panel displays the ADAR knockdown reads. In **b**, orange marks correspond to A->G mismatches and in **a, c**, and **d**, positions marked with blue mismatches are T->C mismatches (A->G on the negative strand).

### Long reads clarify the transcriptional context of inosines

Previous studies have established a connection between editing and changes in splicing, either in *cis* or *trans* [14]. However, we were not able to find many convincing cases of alternative splicing from ADAR knockdown alone with the Illumina data. We ran the differential splicing analysis tools juncBASE [57] and JUM [58] (see Methods). None of the identified splicing events was significant after multiple testing corrections. With our nanopore data, we sought to find edits associated with the presence of other edits or splicing changes that could be overlooked in the Illumina data due to potential mapping difficulties or length limitations. We performed a systematic analysis of all inosine-inosine associations within single molecule reads. For each inosine, we looked at the nearest 20 variants, checked all of the reads that overlapped both variants to count the frequency they co-occured with each other, and performed a Fisher’s test to discover significantly associated positions. We observed 12 associated inosines that satisfied these conditions with a Fisher’s exact p-value < 0.05. In *MRPL30*, we noted coordinated inosine editing occurring more than 500 bp apart within Alu elements (p=2.35e-6) (Fig 4b). The predicted secondary structure of the 3’ UTR consists of a hairpin bringing the two sites in closer proximity. We also noticed a pattern in the 3’ UTR of melanoregulin (*MREG*) transcripts whereby splicing alterations appeared to be coordinated with A-to-I edits. Our nanopore data show that for splicing within the 3’ UTR of *MREG*, there are several positions proximal to splice sites that are edited and unspliced in the CTRL KD samples (Fig 4c). The STAR short-read aligner did not report these splice junctions and there is a lack of Illumina reads aligning across this *MREG* splice site. We then looked for other genes that demonstrated the same mutually exclusive pattern of reads either containing an inosine or having an intron spliced out. We found 145 hyperedited sites that resided within introns of other reads assigned to that gene (Supplementary Data File 2). Three of these sites can be found in the 3’UTR of *CWF19L1* (Fig 4d). These cases illustrate the ways in which long reads can provide benefits over short reads, drawing connections between A-to-I edits where length limitations would affect short read analyses.

## Discussion

The additive complexity of RNA editing and splicing on the transcriptome, in addition to the disease implications of aberrations in these processes, necessitate methods for more thorough profiling of RNA transcripts. We sought to bridge our understanding of A-to-I editing using short- and long-read sequencing to identify edits more extensively as well as investigate any events that require the full transcriptional context to decipher. We knocked down ADAR in lung adenocarcinoma cells and sequenced the cDNA with the more accurate R2C2 nanopore sequencing method. We were able to discover novel type I and type II hyperediting (Fig 2e and 3a), sites that are coordinated with each other (Fig 4b), and sites that may disrupt splicing (Fig 4c,d). Previous work in Alzheimer’s disease has found similar patterns of coupled enriched A-to-I editing in isoforms with longer 3’ UTRs [59]. In this study, we found cases where 3’UTRs were spliced or edited in a mutually exclusive manner. From another study, ADAR-dependent editing of the 3’UTR has been observed to increase expression [60]. This suggests that the elevated levels of editing present in H1975 cells could promote expression of those transcripts bearing edits in their 3’UTRs. The documentation of these transcripts that are targets of ADAR and alternative splicing brings attention to further efforts to disentangle these regulatory processes involved in the cancer transcriptome.

Future studies would benefit from selection of longer molecules to sequence incompletely spliced RNAs and thus capture more intronic A-to-I editing. Nevertheless, we were still able to build computational pipelines to leverage our accurate nanopore data in ways that surpassed the limitations of short reads, continuing to pave a way for the adoption of long reads for characterizing RNA splicing and editing in cancers.

## Methods

### Cell culture and siRNA knockdown

H1975 cells were cultured in T75 flasks with DMEM + 10% FBS media. Cells were split 1:4 every 3 days using a 0.25% trypsin 0.52 mM EDTA solution. Trypsin solution was neutralized using an approximately equal volume of media.

For ADAR and control knockdowns, we used one of the Thermo Fisher Silencer Select siRNAs s1007, s1008, and s1009 for the three *ADAR1* biological replicates and Silencer Select Negative Control No. 1 at 15 nmol for 72 hours. Cells that would be subject to RNA extraction were cultured in 10 cm dishes. In tandem, cells were plated for western blotting in 6-well plates. Depending on the vessel media volume, the appropriate amount of siRNA was added to the media when the cells were 80% confluent.

### Western blotting

We have uploaded our Western blotting protocol to protocols.io [61]. Briefly, after siRNA treatment, the media were aspirated off and 200 ul of cold RIPA and proteinase K solution were added to each well. Cells were scraped off, transferred to cold tubes, and centrifuged at 15,000 x g for 7 minutes. Leaving the pellet, the supernatant lysate was then retained in protein lo-bind tubes. Protein lysates were sonicated twice for 30 seconds, with 1 minute on ice in between rounds of sonication. Protein concentrations were measured with the Pierce BCA Protein Assay. According to the concentrations to ensure approximately equivalent amounts of protein, the lysates were loaded into Bio-rad Mini-PROTEAN precast gels. We used ADAR1 primary antibody (Abcam ab88574) and goat secondary (Abcam ab205719) and imaged on LI-COR C-digit blot scanner.

### RNA extraction

Media was aspirated off and the dishes were washed 3x with ice cold dPBS. 1 ml of tri-reagent was added to each dish and cells were scraped off. Cells suspended in tri-reagent were used as input into the Zymo Direct-zol kit. Following elution from the Direct-zol kit, RNA quality and concentration were evaluated with a Nanodrop, Qubit, and Tapestation (RIN 9.5-9.8).

### R2C2 and nanopore sequencing

We followed the R2C2 protocol from Vollmers et. al. [62]. We also have the workflow written out on protocols.io (https://www.protocols.io/view/r2c2-protocol-draft-n2bvjx3knlk5/v1). In summary, our steps were as follows: Each ADAR KD sample was pooled with a control KD sample. Prior to pooling the samples, 200 ng of RNA were reverse transcribed with SmartSeq and barcoded oligo-dTs. For the ADAR KD and control KD samples that were sequenced together as Pools 1 and 2, 1 ug of RNA was used for the RT step. The RT product underwent lambda exonuclease and RNase A digestion, followed by 15 cycles of PCR using KAPA Hifi HotStart ReadyMix. Next, cDNA was cleaned with 0.8:1 ampure bead purification. The samples that were combined into pool 2 were cleaned using Zymo Select-A-Size for fragments larger than 300 nt, adding an extra empty spin step after the second wash. Pool 1 had 4 samples pooled together. Two were size-selected for fragments larger than 3 kb using a low-melt agarose gel extraction. The two samples to be size-selected were first pooled, then run on a 1% low-melt agarose gel made with TAE. A gel slice containing cDNA above 3 kb was cut out and placed in twice the volume of beta-agarase buffer, incubating on ice. The buffer was refreshed after 20 minutes. After another 20 minutes, the buffer was removed and the gel was melted at 65 C for 10 minutes. The gel was then incubated overnight with the addition of 2 ul of beta-agarase per 300 ul of gel. A bead purification was performed on the DNA-containing digested gel. Final R2C2 cDNA concentration was assessed on a Nanodrop prior to nanopore sequencing preparation. 200 ng of nanopore library was loaded onto a flow cell at time. Excess library was stored at 4C. After 24 hours, any remaining library was loaded after flushing the flow cell with Nuclease flush buffer and DNAse I according to ONT protocol. Reads were basecalled with guppy 4.4.1 and consensus called and demultiplexed using C3POa v2.2.3.

### FLAIR2: splice site fidelity checking and novel isoform detection

FLAIR2 was used for this study, using the –annotation_reliant argument in the FLAIR *collapse* module to invoke the additional transcript alignment step for improved isoform detection. The v2 collapse module first performs an ungapped alignment of reads to transcripts. The top transcript alignments for each read, as determined by minimap2 [63] mapping quality score, are examined with increased stringency: we used the –stringent and –check_splice parameters in FLAIR *collapse* to improve accuracy of read-isoform assignments particularly around splice sites. The --stringent parameter enforces that 80% of bases match between the read and assigned isoform as well as that the read spans into 25 bp of the first and last exons. The --check_splice parameter enforces that 4 out of 6 bases flanking every splice site in the transcript are matched in a given read and that there are no indels larger than 3 bp at a splice site. These new options are incorporated in FLAIR v.1.5.1 and above; subsequent improvements have been made to test the code and make it more user-friendly.

Reads that match annotated isoforms in accordance with these parameters are attached to the isoform as a supporting read. Isoforms with enough supporting reads are included in the final set of FLAIR isoforms. The remaining, unassigned reads are used for novel isoform detection. The process of summarizing the unassigned reads into the isoforms begins with minimap2 to align the reads to the genome. FLAIR corrects unsupported splice sites with the closest splice site that contains more evidence i.e. splice sites found in annotations or short-read sequencing. The corrected reads are then grouped by their splice junction chains. For each group, FLAIR calls transcription start and end sites, collapsing each group into one or more representative first-pass isoform. Next, FLAIR assigns each read to a first-pass isoform by realigning the reads to the isoforms and identifying the best alignment with the splice site fidelity stringency previously specified. The final FLAIR isoform set arises from filtering the first-pass set for the novel isoforms that pass a minimum supporting read threshold combined with the annotated isoforms.

### SIRV analysis

We analyzed SIRV reads that aligned with the SIRV1-SIRV7 references from the LRGASP mouse embryonic stem cell R2C2 sequencing replicates [39]. We ran FLAIR2 providing the genome annotation and with the default minimum supporting read count of 3 (-s). We used the -L parameter and supplied a genome annotation for the stringtie2 v2.0 run, using the default parameters otherwise. For FLAMES (cloned from GitHub June 1, 2021), we used the provided SIRV config file and altered the minimum supporting read count of 3. We used gffcompare [64] to calculate transcript-level sensitivity and precision of each tool’s transcript reference with the ground truth, using a wiggle room of 50 bp at the transcription start sites and terminal ends for matching (-e 50 and -d 50).

### Variant integration into FLAIR isoforms

We ran two long-read variant callers on our data. Longshot was run with default and required arguments in addition to --min_allele_qual set to 3 and the -F argument. Pepper-Margin-DeepVariant version r0.7 was run on a bam file with all cigar string N operators changed to D and H operators removed. We ran Pepper-Margin-DeepVariant with default and required arguments including the --ont_r9_guppy5_sup argument. For the longshot-phased version of FLAIR, we supplied FLAIR-collapse the longshot bam and vcf output using the --longshot_bam and -- longshot_vcf arguments, respectively. For the variant caller-agnostic method, we filtered the vcfs generated from longshot and pepper by coverage and then combined them. This vcf, along with FLAIR-collapse isoform output, was supplied to a FLAIR script called assign_variants_to_isoforms. Both methods of variant integration ultimately yield a sequence fasta file with the variant-containing isoform sequences, an updated isoform model file, as well as a vcf of the variants and the isoform names that contain those variants.

### Illumina RNA-Seq analysis

REDItools was used to tabulate the number of reads supporting each base at every position. The REDItools output was filtered using custom python scripts for positions that contained guanosine mismatches at positions where the reference base was an adenosine for genes corresponding to the forward strand of the genome, and the reverse complement for those on the reverse strand. Positions with less than 15% putative editing were filtered out. The counts of the reference and alternate allele in each of the samples for the remaining positions were supplied to DRIMSeq [65] for differential testing between two conditions, with the settings that at least 5 reads contained editing (G mismatch) in a minimum of two samples, as well as a coverage of 15 reads minimum in at least 3 samples. We ran juncBASE v1.2-beta following the manual with default parameters. We ran jum v2.0.2 with the parameters ‘--JuncThreshold 5 --Condition1_fileNum_threshold 2 -- Condition2_fileNum_threshold 2‘ and default parameters otherwise.

### Inosine detection in long reads

We used the pysam python package’s pileup method to count A->G or T->C reads at variant positions. Next, we combined our nanopore data by knockdown condition, followed by filtering for positions that had a minimum coverage of 10 in either condition and a change in percentage of edited reads after ADAR knockdown of 10% or more. We performed a Fisher’s exact test to assess the significance of the A-to-I differences.

### Inosine coordination analysis for long reads

For the long-range inosine coordination analysis, we considered inosines that were at least 50 bp apart. We looked at the secondary structure of *MPRL30* by inputting its 3’ UTR sequence to the RNAfold web server. For the inosine-intron coordination analysis, we filtered for sites that were type I hyperedited (i.e. more than 40% of residues were edited) and had at least 10 reads that were edited. We also required that at least 10 reads had the edited position spliced out.

## Code Availability

FLAIRv2.0 is available on GitHub at https://github.com/BrooksLabUCSC/flair.

## Data Availability

The R2C2 consensus read sequences have been submitted to SRA under bioproject PRJNA981664. FLAIR2 isoforms with longshot variants have been uploaded to Zenodo at https://zenodo.org/record/8019704.

## Author Contributions

A.D.T. and A.N.B. designed the study. E.H.R., R.V., and C.V. assisted in the design of experiments. A.D.T. performed experiments. A.D.T. wrote code and analyzed data. R.V. and C.V. processed data. A.D.T. and A.N.B. interpreted the data. A.D.T. and A.N.B. wrote the manuscript.

## Acknowledgments

We would like to acknowledge Sergio Covarrubias, Apple Vollmers, and Mays Mohammed Salih for guidance with ADAR knockdown. We also would like to acknowledge Dori Zhiqian Deng, Alexander Zee, and Matthew Adams for guidance on R2C2 library preparation. This work was supported by NIGMS R35GM138122 (A.N.B.) and F31HG010999 (A.D.T.).

---

**Supplementary Table 1. HST-containing genes in Castaneus x Mouse 129 identified from this study.**

**Supplementary Table 2. GO terms associated with haplotype-specific transcripts found in hybrid castaneus x 129 mouse data.**

**Supplementary Table 3. DESeq2 output table of differentially expressed genes upon KD of ADAR in H1975.** Analysis is from Illumina sequencing data.

**Supplementary Data File 1. Significant A-to-I changes identified in R2C2 nanopore data**. Columns are as follows: 1) chromosome and 2) position of inosines, the number of reads in the control KD sample that are 3) not edited or 4) edited, the number of reads in the ADAR KD samples that are 5) not edited or 6) edited, and the 7) Fisher’s exact test p-value.

**Supplementary Data File 2. A-to-I editing within intronic regions**. Columns are as follows: 1) chromosome and 2) position of inosines, the number of 3) unedited reads, 4) edited reads, and 5) number of reads with the inosine spliced out.

**Supplementary Figure. 1.**
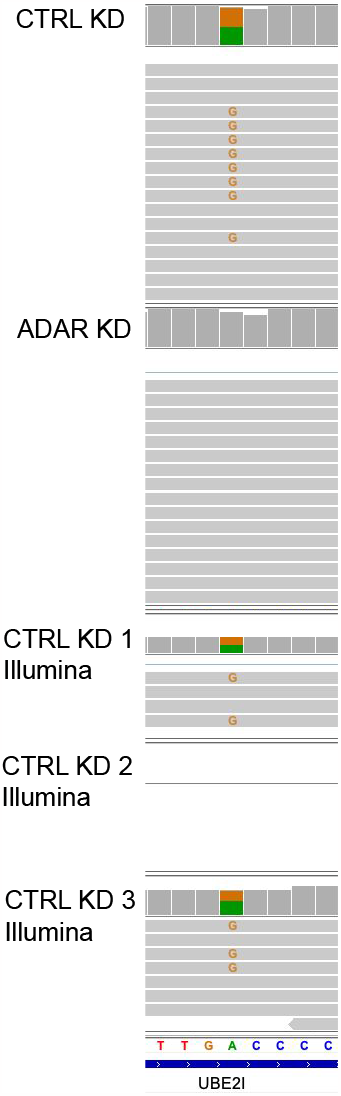
Example of novel A-to-I editing found with nanopore data. IGV shots of nanopore and Illumina data aligned to hg38. There were no reads aligning to *UBE2I* in the second Illumina CTRL KD replicate.

**Supplementary Table. 2.** GO terms associated with haplotype-specific transcripts found in hybrid castaneus x 129 mouse data.

| Gene Set Name | p-value | FDR q-value |
| --- | --- | --- |
| DNA metabolic process | 2.75E-13 | 4.27E-09 |
| DNA repair | 2.21E-11 | 1.72E-07 |
| Cellular response to DNA damage stimulus | 1.80E-10 | 9.34E-07 |
| Cellular response to stress | 3.39E-10 | 1.32E-06 |
| Chromosome | 1.45E-09 | 4.49E-06 |
| Cell cycle | 3.27E-09 | 8.47E-06 |
| Transferase complex | 7.38E-09 | 1.62E-05 |
| Catalytic complex | 8.34E-09 | 1.62E-05 |
| Small molecule metabolic process | 6.89E-08 | 1.19E-04 |
| Cellular macromolecule biosynthetic process | 1.72E-07 | 2.67E-04 |

